# Phage display uncovers a sequence motif that drives polypeptide binding to a conserved regulatory exosite of O-GlcNAc transferase

**DOI:** 10.1101/2023.03.15.532872

**Authors:** Matthew G. Alteen, Richard W. Meek, Subramania Kolappan, Jil A. Busmann, Jessica Cao, Zoe O’Gara, Ratmir Derda, Gideon J. Davies, David J. Vocadlo

**Author notes:** Corresponding Author: David J. Vocadlo. **Author Contributions:** M.G.A., R.W.M., R.D., G.J.D., and D.J.V. designed research, M.G.A., R.W.M., S.K., J.A.B. performed research. M.G.A., R.W.M., S.K., J.A.B., J.C., Z.O., R.D., G.J.D., and D.J.V. analyzed data, M.A. and D.J.V. wrote the paper with input from the other authors. **Competing Interest Statement:** J.C., Z.O, and R.D. are employees or consultants with 48 Hour Discovery.

## Abstract

The modification of nucleocytoplasmic proteins by O-linked N-acetylglucosamine (*O*-GlcNAc) is an important regulator of cell physiology. *O*-GlcNAc is installed on over a thousand proteins by just one enzyme, *O*-GlcNAc transferase (OGT). How OGT is therefore regulated is therefore a topic of interest. To gain insight into these questions, we used OGT to perform phage display selection from an unbiased library of ∼10^8^ peptides of 15 amino acids in length. Following rounds of selection and deep mutational panning we identified a high-fidelity peptide consensus sequence, [Y/F]-x-P-x-Y-x-[I/M/F], that drives peptide binding to OGT. Peptides containing this sequence bind to OGT in the high nanomolar to low micromolar range and inhibit OGT in a non-competitive manner with low micromolar potencies. X-ray structural analyses of OGT in complex with a peptide containing this motif surprisingly revealed binding to an exosite proximal to the active site of OGT. This structure defines the detailed molecular basis driving peptide binding and explains the need for specific residues within the sequence motif. Analysis of the human proteome revealed this motif within 52 nuclear and cytoplasmic proteins. Collectively, these data suggest an unprecedented mode of regulation of OGT by which polypeptides can bind to this exosite to cause allosteric inhibition of OGT through steric occlusion of its active site. We expect these insights will drive improved understanding of the regulation of OGT within cells and enable the development of new chemical tools to exert fine control over OGT activity.

**SIGNIFICANCE STATEMENT:** Thousands of proteins within humans are modified by the monosaccharide N-acetylglucosamine (O-GlcNAc). O-GlcNAc regulates cellular physiology and is being pursued to create therapeutics. Remarkably, only one enzyme, O-GlcNAc transferase (OGT), installs O-GlcNAc and its regulation is poorly understood. By affinity selection using a vast peptide library, we uncover an amino acid sequence motif that drives binding of polypeptides to OGT. An OGT-peptide complex shows how this motif binds to an allosteric site proximal to the active site and inhibits OGT in an unprecedented manner. Given the distribution of this sequence motif within the human proteome proteins containing this motif likely regulate the activity of OGT, outlining a new mode by which OGT is controlled and opening new avenues for research.

## INTRODUCTION

Post-translational glycosylation of proteins has emerged as serving essential roles in regulating cellular functions.(1) One form of glycosylation involves addition of *N*-acetylglucosamine (GlcNAc) to the hydroxyl groups of serine and threonine residues of nuclear and cytosolic proteins(2, 3). This modification, known as *O*-GlcNAc, modulates human physiology through its diverse molecular roles in regulating transcription,(4, 5) protein stability,(6) and stress response(7). Despite the widespread occurrence of this post-translational modification on hundreds of protein targets,(8) cycling of *O*-GlcNAc is controlled by just two enzymes. The glycosyltransferase (GT) *O*-GlcNAc transferase (OGT) from family 41 (GT41) catalyzes the installation of *O*-GlcNAc to acceptor polypeptides using uridine diphosphate GlcNAc (UDP-GlcNAc) as the glycosyl donor.(9) Hydrolytic cleavage of *O*-GlcNAc is effected by the glycoside hydrolase (GH) from family 84 (GH84) known as *O*-GlcNAcase (OGA), which hydrolyzes the glycosidic linkage to liberate free serine or threonine residues.(9, 10) Since dysregulation of *O*-GlcNAc cycling has been genetically linked to X-linked Intellectual Disability (XLID)(11, 12) and also implicated in numerous pathological conditions, including neurodegenerative diseases(13, 14) and cancers,(15, 16) there is considerable interest in studying the regulation of these two enzymes and generating new tools and insights that can be exploited to enable their manipulation.(17)

The structure of OGT consists of two principal parts: an N-terminal superhelical section comprising 13.5 tetratricopeptide repeat (TPR) domains and a C-terminal catalytic region (**Figure 1a**).(18-20) This C-terminal catalytic region can be divided into three folded domains; two catalytic domains (N-Cat and C-Cat) that together comprise the active site and an intervening domain (Int-D) which separates the catalytic domains and has a yet undefined function.(9, 21) Selection of protein substrates by OGT is thought to be mediated by its TPR domains,(21, 22) which form a superhelical groove extending into the active site of OGT (**Figure 1b** and **1c**), but molecular details as to how it does this remain ill defined. An asparagine ladder lining the TPR superhelical groove has been found to coordinate the peptide backbone(23) and aspartate residues enables selection of a protein subset, but how OGT achieves greater specificity for various sets of proteins remains unknown.

**Figure 1.**
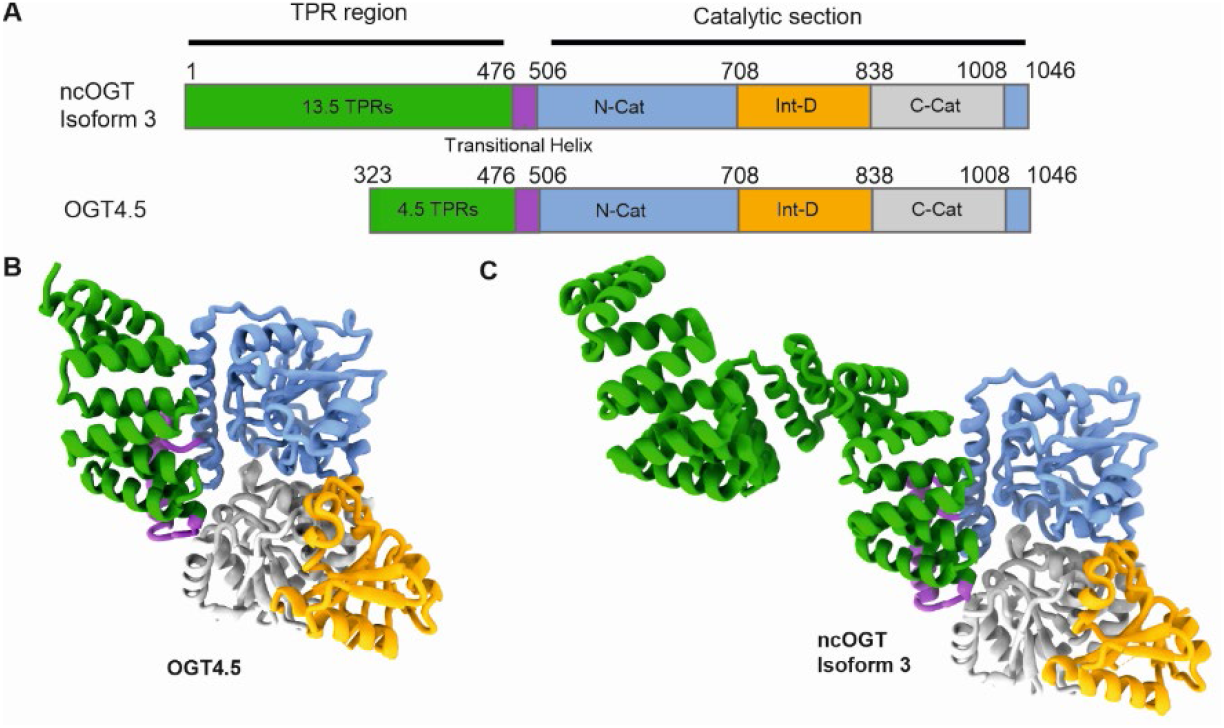
Domain arrangement of OGT: (A) Schematic showing the distribution of folded domains across the primary sequence of full-length OGT (ncOGT) and the crystallized form of OGT (OGT4.5). Tetratricopeptide repeats (TPRs), N-terminal catalytic domain (N-Cat), intervening domain (Int-D), and C-terminal catalytic domain (C-Cat) are shown with the amino numbers defining the domain boundaries. (B) The domains, coloured as in the primary sequence representation, mapped onto the OGT4.5 crystal structure and (C) onto the structure of full-length ncOGT (isoform 3).

Because OGT has hundreds of known protein targets, yet lacks a clear consensus sequence governing glycosylation, the activity of OGT is thought to be regulated in various ways by cellular factors.(21, 24, 25) Among these regulatory mechanisms, OGT dimer status has been found to influence its activity(20) as has an interaction with an intrinsically disordered region of OGA.(26) Interactions between OGT and other proteins, mediated through direct binding to the TPR region, have also been proposed to regulate OGT activity.(26) This last idea has gained support through quantitative proteomics studies showing that different cellular conditions alter the set of its interacting partners.(27, 28) Interestingly, direct binding of candidate proteins identified in these proteomic studies has not yet been observed and how such proteins might modulate OGT activity remains to be elucidated. In brief, how OGT is regulated remains enigmatic and a topic of considerable interest.

Among various approaches to aiding understanding of the roles of OGT is the use of inhibitors of this enzyme.(29, 30) Moreover, OGT inhibitors may hold promise for the treatment of conditions, including various cancers, which are linked to excessive *O*-GlcNAcylation.(15, 16) However, despite several high-throughput screening campaigns to identify small-molecule inhibitors of OGT, most compounds arising from these efforts have limited selectivity and cell-permeability.(31, 32) Extensive medicinal chemistry efforts, however, have led to OSMI-4, a nanomolar inhibitor OGT that shows utility within cells.(33) We noted that identifying these HTS-derived inhibitors relies on activity-based assays or competitive displacement assays that are biased towards the discovery of compounds binding to the active site. Accordingly, compounds that bind OGT at alternative sites could be useful complements to the existing set of competitive OGT inhibitors. Toward this goal of identifying ligands that bind to OGT, we recently turned to display methods as an alternative approach to uncover ligands for OGT.(34) We used the RaPID mRNA display system(35) to identify cyclic peptide inhibitors of OGT that bind to the TPR domain and exert allosteric inhibition.(34, 36) In parallel to those efforts, we turned to using standard phage display as an alterative approach for the discovery of linear peptides that bind OGT. We expected that the advantages of DNA-encoded linear peptide libraries, including high structural diversity, synthetic accessibility, and general biocompatibility, could uncover peptide sequences binding to the TPR domains or active site that could serve as interesting leads for new inhibitors.(37, 38) Additionally, since phage display screening methods rely on affinity-based selection, we reasoned that we might identify peptide ligands of OGT found within native proteins that may be either inhibitors or simply non-inhibitory ligands. In pursuit of this goal, we screened immobilized OGT against a deep library comprising ∼10^8^ sequences of linear 15-amino acid peptides expressed on the surface of the pIII major coat protein of filamentous M13 phage. After identification of sequence preferences and affinity maturation of the selected phage, a panel of peptides with submicromolar non-competitive inhibition of OGT were discovered. Furthermore, we made the surprising observation that these peptides bind an exosite within the intervening domain of the C-terminal section of OGT. Finally, we showed that inhibition by these peptides is correlated with the length of their sequences, consistent with a steric occlusion model that antagonizes binding of polypeptide substrates to the active site of OGT.

## RESULTS

### Phage library development and screening results

To identify potential peptide ligands of OGT, we screened full length OGT against an unbiased phage-display library constructed using codon-corrected trinucleotide phosphoramidite cassettes (TriNuc), which produce a single codon for each amino acid and result in equal distribution of amino acids within translated sequences.(39) This library strategy eliminates amino acid sequence representation biases arising from variable codon redundancies within the standard genetic code. As a first step, we used a TriNuc-encoded library containing 19 codons for each amino acid, excluding cysteine, to produce a library of linear peptides comprising 15 randomized amino acids. These peptides were presented on M13 phage by fusion to the *C*-terminus of the pIII coat protein. This resulted in a library we termed X15 that has a sequence diversity of ∼10^8^. We immobilized recombinant full-length OGT, bearing a hexahistidine-tag (His_6_) fused to the *N*-terminus, on magnetic nickel-NTA beads and then incubated the beads with the X15 phage library (**Figure 2a**). We performed four rounds of phage panning to enrich phage binding to OGT from the larger phage population and analyzed enriched sequences from selected samples (**Figure 2b** and **2c, Table S1**) by lysis followed by DNA isolation and next-generation sequencing (NGS) (**Figure S1**).(40)

**Figure 2.**
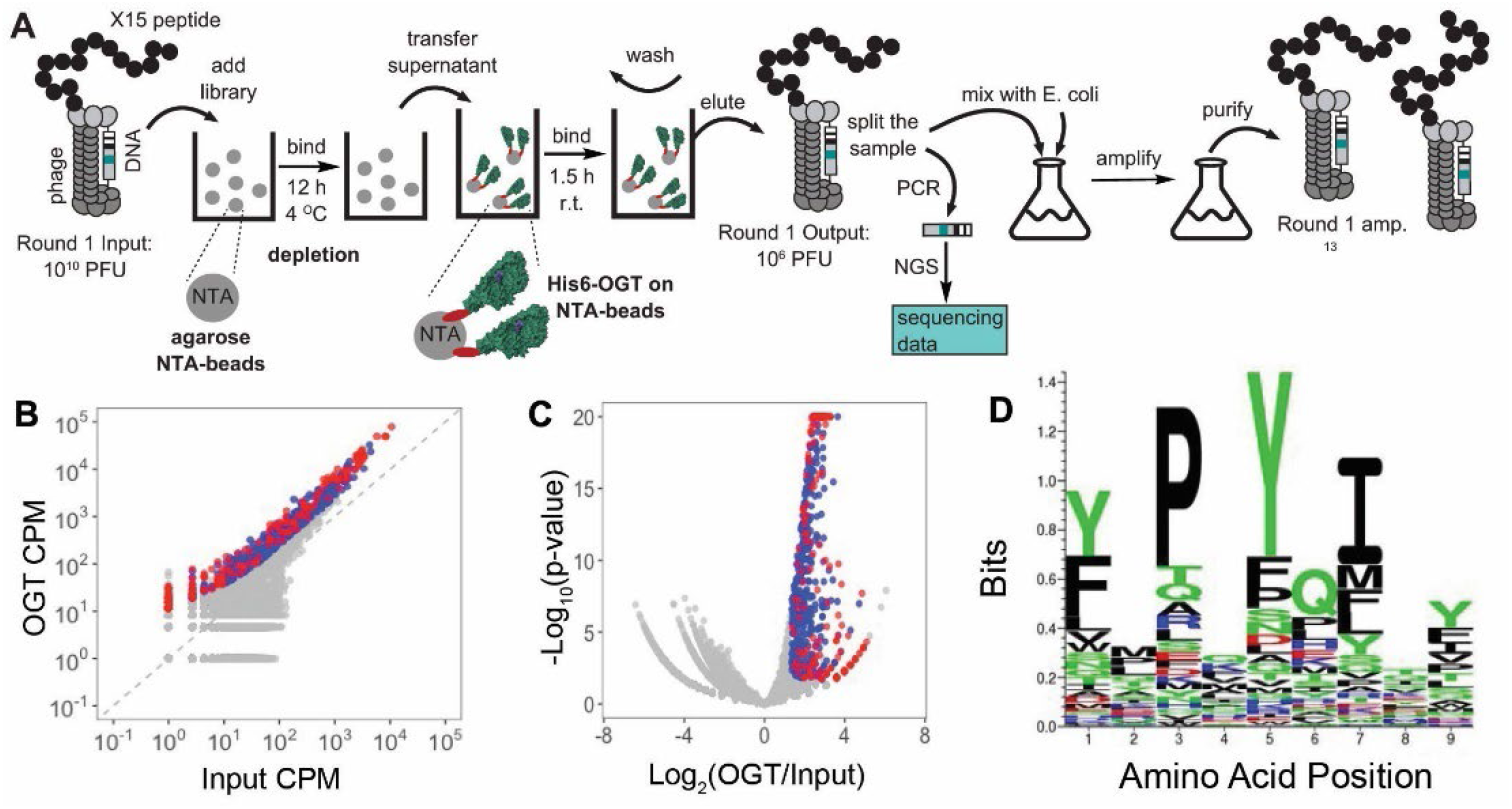
Outline of phage display screening approach and sequence conservation of the peptide binders of OGT: (A) Schematic of one round of the phage panning procedure using bead-immobilized OGT. In each round, the phage library was depleted against Ni-NTA beads to generate the input library, which was incubated with bead-immobilized OGT, followed by washing and elution. Samples from the resulting phage library were submitted for next generation sequencing (NGS), and also amplified in *E. coli* for the next round screening. (B-C) Bioinformatic analysis of Round 3 of screening NGS data. Each dot represents a sequence that observed in the NGS data, coloured dots are 645 sequences that pass differential enrichment (DE) analysis and hence exhibit significant enrichment in OGT samples whereas grey dots have not passed the DE test. The left scatter plot (B) shows the counts per million (CPM) of each sequence in bound to bead-immobilized OGT compared to Input whereas the right plot shows comparison of the same Input in binding to NiNTA-magnetic beads. (D) Sequence logo indicating the prevalence of amino acid residues in top 300 peptide binders to OGT after four rounds of phage panning.

Analysis of the sequencing results was accomplished using multiple strategies. Initially, we ranked the sequences identified through NGS according to their normalized sequence count per million sequences (NCPM). Notably, out of 18,000 total sequences in the final enriched phage library, 81% of the overall enrichment as assessed using NCPM values were attributable to just the top 200 sequences (**Table S1**), revealing a high degree of enrichment for this set of peptides. In round 3,645 sequences also passed a differential enrichment analysis (DE) in which the sequences were significantly enriched in binding to OGT when compared to input and but not in the experiments that used protein-free bead control. By the fourth round of phage panning the enrichment with respect to input was modest (**Figure S1e**) and we therefore stopped selections at this point. We then binned the sequences into groups of 100 that we analyzed using the Kalign2 multiple sequence alignment tool.^23^ This analysis revealed that among these top 200 sequences a strong consensus motif was present with 90% fidelity. Further analysis uncovered that the motif, [Y/F]-x-P-x-Y-x-[I/M/L] dominated the top enriched sequences in rounds 3 and 4 (**Figure 2d**) and it was detectable as early as round 2 of panning (**Figure S1**). Revealingly, this seven-amino acid motif was present at different registers within the 15-amino acid sequence of the displayed peptide, indicating that variability at either end of the motif did not markedly impair binding to OGT. We observed some variability in positions between the conserved residues of this motif, suggesting oriented binding of these peptides to OGT. We also remarked on the presence of a conserved proline residue within this sequence that is consistent with observations from large-scale proteomic studies on the peptide substrate preferences of OGT. This last observation suggested to us that these peptides may bind within the active site of OGT.(8, 41, 42)

### In vitro inhibition of OGT by prioritized peptide sequences

Having identified putative binders, we next proceeded to use synthetic peptides to examine whether these candidate ligands might bind to OGT. As noted above, the proline within the observed consensus motif suggested to us that these peptides were likely binding within the active site, which pointed to inhibition experiments as being potentially informative. We prioritized peptides selected from among the top 30 enriched sequences as assessed by NCPM. All these sequences exhibited at least a 1000-fold enrichment over the level seen for background peptide sequences (**Table S1**). The top 30 sequences were then grouped into five families based on amino acid composition within the consensus region and a total of 10 peptides, two from each family, were synthesized for testing (**Table 1**). We then assessed whether these 10 peptides could inhibit the glycosyltransferase activity of OGT using a direct fluorescence activity-based OGT assay.(43) Using this assay, we first screened these peptides for inhibition of OGT at a concentration of 100 μM and identified four peptides from four families that showed notable inhibition (**Table 1, Figure S2**), while peptide P6 had inadequate solubility for testing. We then generated concentration-response curves for members from these four families and determined their IC_50_ values against OGT, which ranged from 21 to 37 μM (**Figure 3a**). Interestingly, five peptides showed no significant inhibition towards OGT. Because these sequences were likely OGT ligands, this observation suggested that there are features within these peptides, outside of the motif, that are required for inhibition.

**Table 1:**
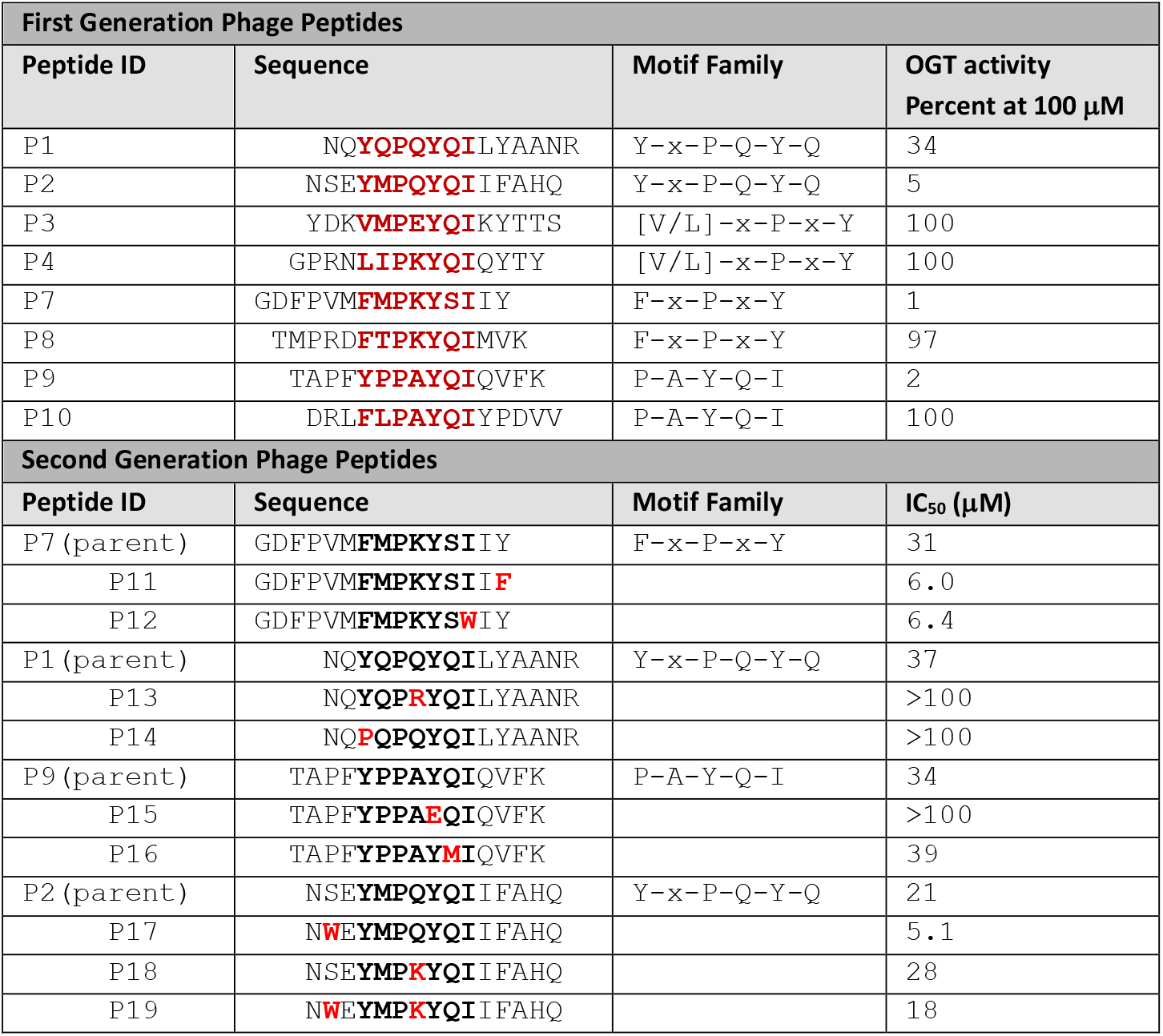
Sequences and OGT inhibition of prioritized peptides from phage panning showing in bold the conserved motifs.

**Figure 3.**
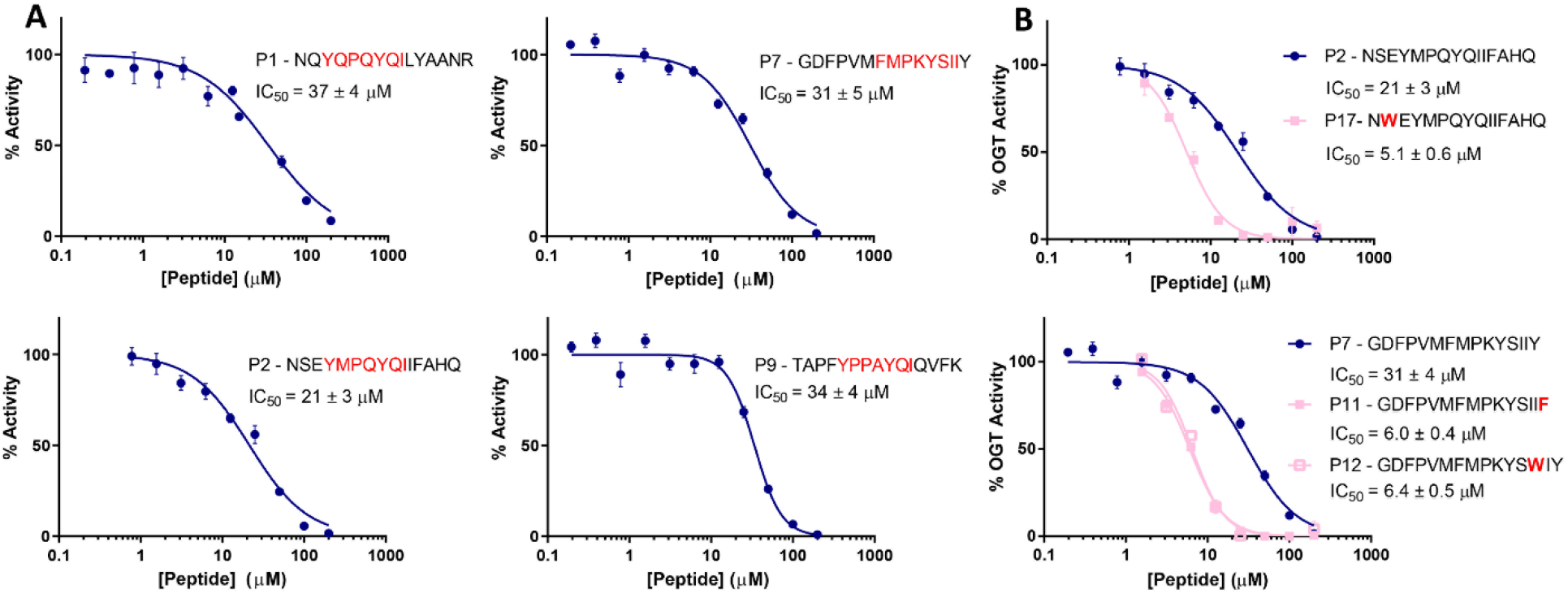
Dose-dependent inhibition of OGT activity by phage display-derived peptides: (A) Concentration-response curves of four of the top peptide binders derived from the first round of phage panning that inhibit OGT. The consensus motif determined from analysis of phage panning results is shown in red for each peptide. % OGT activity was normalized relative to controls containing 1% v/v DMSO (100% activity) or 1 mM UDP (0% activity). All assays were performed in buffer containing 0.02% Triton X-100 detergent. Sigmoidal curves were fit using a four-parameter log(inhibitor) vs response formula in GraphPad Prism 8. Error bars represent S.E.M. of quadruplicate measurements. (B) IC_50_ values for three inhibitory peptides derived from the second round of phage panning against OGT. Inhibitory activities of peptides having lower IC_50_ values shown in comparison with the IC_50_ values obtained for the original sequences identified in the first rounds of selection. Mutations arising in the sequences from round two of the phage selection are in red bolded font.

### Phage library maturation and evaluation of second-generation peptides

After determining IC_50_ values of these peptides from the initial phage panning and thereby validating the importance of this seven-amino acid consensus motif, we next sought to build upon these observations by generating a focused library for subsequent phage selection experiments. We therefore constructed a phage library consisting of 11 amino acids centred on the top four inhibitory sequences, with an additional 762 randomized sequences also included to ensure quality control of the library. We elected to use the 11-mer peptides because this length allowed full positional maturation through saturation mutagenesis. Using this focused phage library, we carried out a new round of phage panning against OGT to identify matured peptide sequences having greater affinity for OGT. While the enrichment of 11-mers appeared to be weaker that 15-mers overall, we were encouraged to discover, after sequencing analyses, that four families of peptides were enriched compared to the overall 11-mer population We ranked these peptides based on their relative fold-enrichment over the parent sequences (**Table S1**) and, on this basis, prepared 10 additional 15mer peptides (**Table 1**), containing overrepresented point mutations. We found three proposed positional substitutions in 15-mer sequences gave rise to peptides with greater inhibitory activity toward OGT than parental 15-mer sequences (**Figure 3b**) with IC_50_ values ranging from 5 to 7 μM, representing a 5-to 6-fold improvement over the corresponding first generation of parental peptides (**Table 1**). Seven other substitutions had no effect or even weaker inhibitory potency than the corresponding parent 15-mer peptide. These observations, including the improvements in inhibitory potency, suggest that while the consensus motif is critical for binding to OGT, the flanking positions and positions between the conserved residues also play a role in the ability of these peptides to inhibit OGT, with hydrophobic residues enhancing binding and hydrophilic residues within or adjacent to the motif being unfavorable.

To better understand how these peptides were inhibiting OGT, we performed kinetic analyses on two of the most potent affinity matured peptides (P11 and P17) to identify their mode of inhibition (**Figure 4a** and **4b, Figure S3**). We accomplished this by varying either the concentration of the fluorescent UDP-GlcNAc donor or the peptide acceptor used in the OGT activity assay in the presence of several concentrations of peptide OGT-P17. Surprisingly, global fitting of the data revealed both P11 (**Figure 4a** and **4b**) and P17 (**Figure S3**) inhibited OGT in a non-competitive manner with respect to the acceptor peptide and donor substrate. The calculated *K*_I_ value for P11 inhibition with respect to UDP-GlcN-BODIPY was 2.7 ± 0.3 μM and with respect to Biotin-HCF was 3.8 ± 0.4 μM (R^2^ = 0.87 and 0.85). The non-competitive mode of inhibition, coupled with the concordance in *K*_I_ values for P11 inhibition with respect to each substrate, strongly suggested that binding of the peptides occurs outside of the active site of OGT. Given the peculiar mode of inhibition of P11, we decided to determine the *K*_d_ of P11 using biolayer interferometry (BLI). Immobilization of biotinylated P11 onto the chip surface followed by monitoring of binding by OGT showed binding reached equilibrium over time (**Figure 4c**). Plotting these equilibrium positions reflected by the maximum response (R_max_) and revealed an excellent fit to a single site – no background binding model that reflected high affinity binding (*K*_d_ = 470 ± 160 nM, R^2^ = 0.99) to OGT (**Figure 4d**). The tighter binding, as reflected by 10-fold lower *K*_d_ value, as compared to inhibition, reflected by the *K*_I_ value, was unexpected but we reasoned it could arise an allosteric mode of inhibition in which a perturbation arising from binding is incompletely transmitted to the active site. These data collectively support that these peptides are binding to OGT and inducing inhibition through an allosteric mechanism, perhaps by inducing a conformational shift that causes inhibition in a manner akin to that observed for recently described macrocyclic peptide inhibitors of OGT.(36)

**Figure 4.**
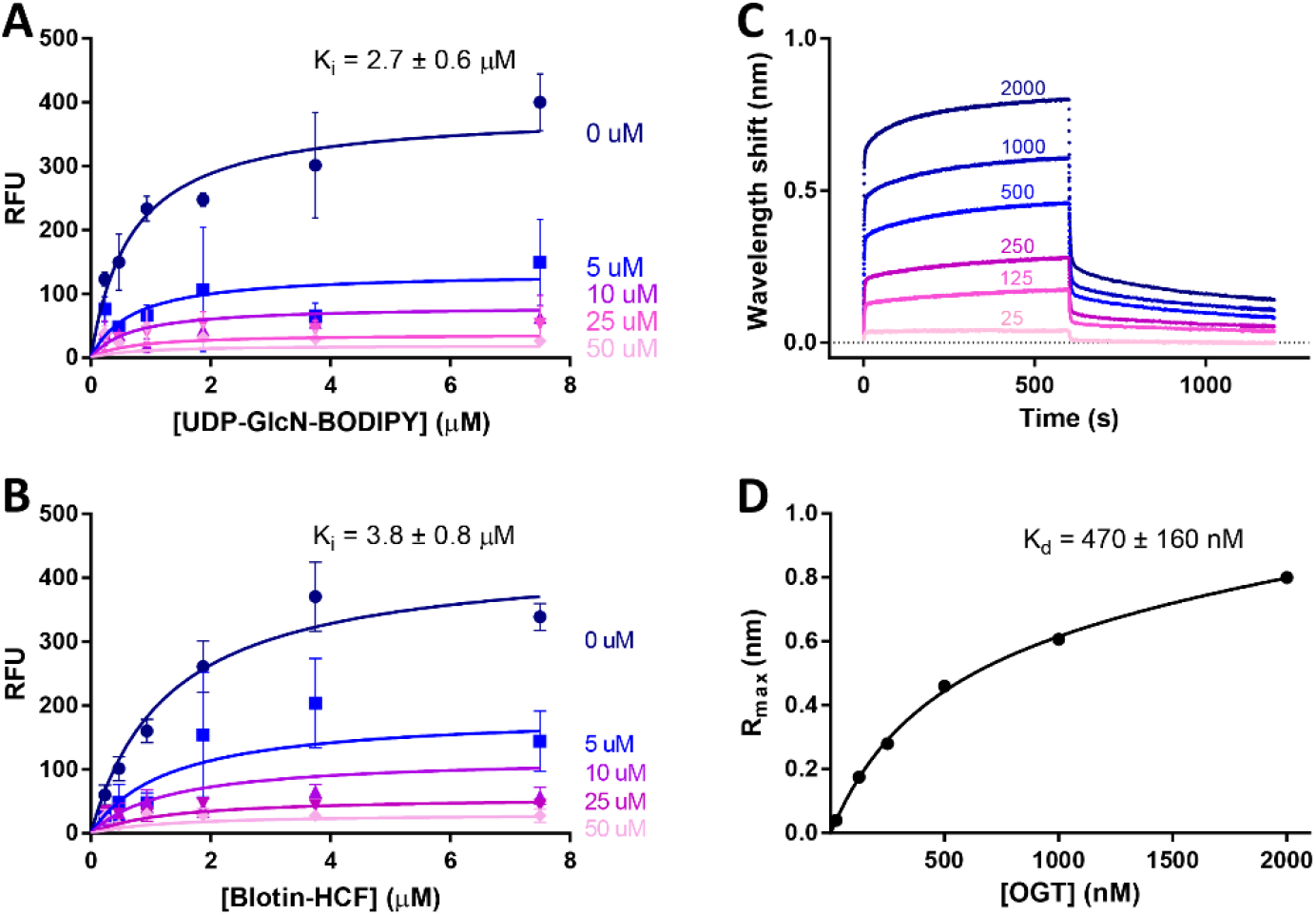
Inhibition and binding of OGT by peptide P11: (A) Inhibition of OGT by P11 at varying concentrations of P11 and UDP-GlcN-BODIPY reveal a mode of non-competitive inhibition. (B) Inhibition of OGT by P11 at varying concentrations of P11 and Biotin-HCF peptide reveal a mode of non-competitive inhibition. (C) Biolayer interferometry data for binding of P11 to OGT determined by immobilization of P11 followed by binding of OGT at different concentrations. (D) Fitting to a single binding model to a plot of maximum response (R_max_) as a function of [OGT] reveals a dissociation constant (*K*_d_) consistent with the *K*_I_ value determined by global fitting of the inhibition data.

### Structure-activity relationships using targeted peptide truncations and deletions

To gain a better understanding of the features of these peptides that mediate inhibition of OGT, we proceeded to assess the inhibitory potency of two truncated peptides (tP11 and tP17) that comprised only the 7-residue consensus motif ([Y/F]-x-P-x-Y-x-[I/M/F]) identified from our sequencing results. Strikingly, none of the short peptides that we tested inhibited OGT activity at concentrations of up to 100 μM (**Table S2, Figure S4**). Such results were also in accordance with overall weak binding potency of shorter 11-mer peptides in a maturation screen. Nevertheless, because the phage screening results contained longer, 15-mer peptides with variable residues beyond the sequence motif, we reasoned that inhibition required the presence of amino acid residues pendent to the consensus motif. We therefore hypothesized that these extensions likely somehow drive inhibition. To investigate this idea, we prepared a series of peptide constructs based around the sequence of the P11 and P17 peptides and then tested these as inhibitors of OGT (**Table S2** and **Figure S4**). As a first step, we enhanced the solubility of P11 to determine whether inhibition was influenced by solubility or aggregation of these peptides. We found that the addition of two lysine residues at either the N- or C-terminus of the peptides (P21 and P22) had only marginal effects on inhibition, indicating inhibition was likely mediated by the soluble monomeric peptide. In addition, deletion of three N-terminal residues (P24) also did not significantly perturb inhibition while deletion of all residues N-terminal of the motif had a modest effect (P25). Conversely, deletion of the two C-terminal residues (P23) resulted in complete loss of inhibition at concentrations up to 200 μM. No inhibition from a control peptide (P26) was observed. These data collectively suggested that, in addition to the importance of the motif, residues outside of the motif at the C-terminal end are essential for inhibition of OGT. In contrast, residues at the N-terminus are less important for either binding or inhibition. Together, these collective data support these peptides authentically binding to OGT and inhibition being sensitive to small changes within the sequence motif, its intervening residues, and the presence of residues C-terminal to the motif.

### X-ray structure of OGT in complex with truncated peptides

Based on the unique binding and inhibition properties of these peptides, coupled with their non-competitive mode of inhibition, we judged that structural insights into their binding to OGT could be illuminating. We therefore set out to determine the X-ray crystallographic structure of OGT in complex with P11. To do so, we used a previously reported, readily crystallizable construct of OGT (OGT4.5) that consists of the catalytic region appended to 4.5 TPRs.(19) To aid crystallisation we used a truncated version of the P11 peptide (tP11; FMPKYSI) bearing just the motif of interest. The residues of peptide tP11 are numbered to match P11. Due to limited peptide solubility, we found it necessary to concentrate OGT4.5 in the presence of the tP11 peptide to obtain a soluble OGT-peptide complex. Crystal screening yielded diffraction quality crystals that were processed in the *P*321 spacegroup to 2.8 Å with four copies of OGT comprising the asymmetric unit (**Table S3**). Most surprisingly, clear *F*_*o*_-*F*_*c*_ electron density, incompatible with the OGT main chain, was observed across a face of the intervening domains (Int-D) (**Figure S5**), suggesting these linear peptides were not binding to the TPR domains as we had expected. Into this density it was possible to unambiguously model the tP11 peptide. The tP11 peptide binds across the Int-D of the catalytic region, a region we hereafter refer to as an exosite, lying outside of the active site (**Figure 5a**). Despite numerous structural insights into OGT, no function has yet been attributed to the Int-D and only OGT enzymes from metazoans possess this domain.(9, 21) While the electron density of tP11 is of sufficient quality to model in the peptide, electron density is diffuse for the sidechains of F7 and I13 (**Figure 5b**). The peptide is bound in place by hydrogen bonding between N791 and the I13 backbone and hydrogen bonding between Y11 and the carbonyl group of L837. In addition, the carbonyl of S833 hydrogen bonds to the backbone nitrogen of K10. (**Figure 5c**). Further interactions occur along the length of the peptide (**Figure 5d**). Notably, I13 in the in the tP11 peptide sequence faces and binds into a hydrophobic pocket formed by F723, I734 and I787 (**Figure 5d**). Strikingly, the residues making up the exosite that cradles this peptide motif are conserved among mammalian OGT orthologues (**Figure 6** and **Figure S6**).

**Figure 5.**
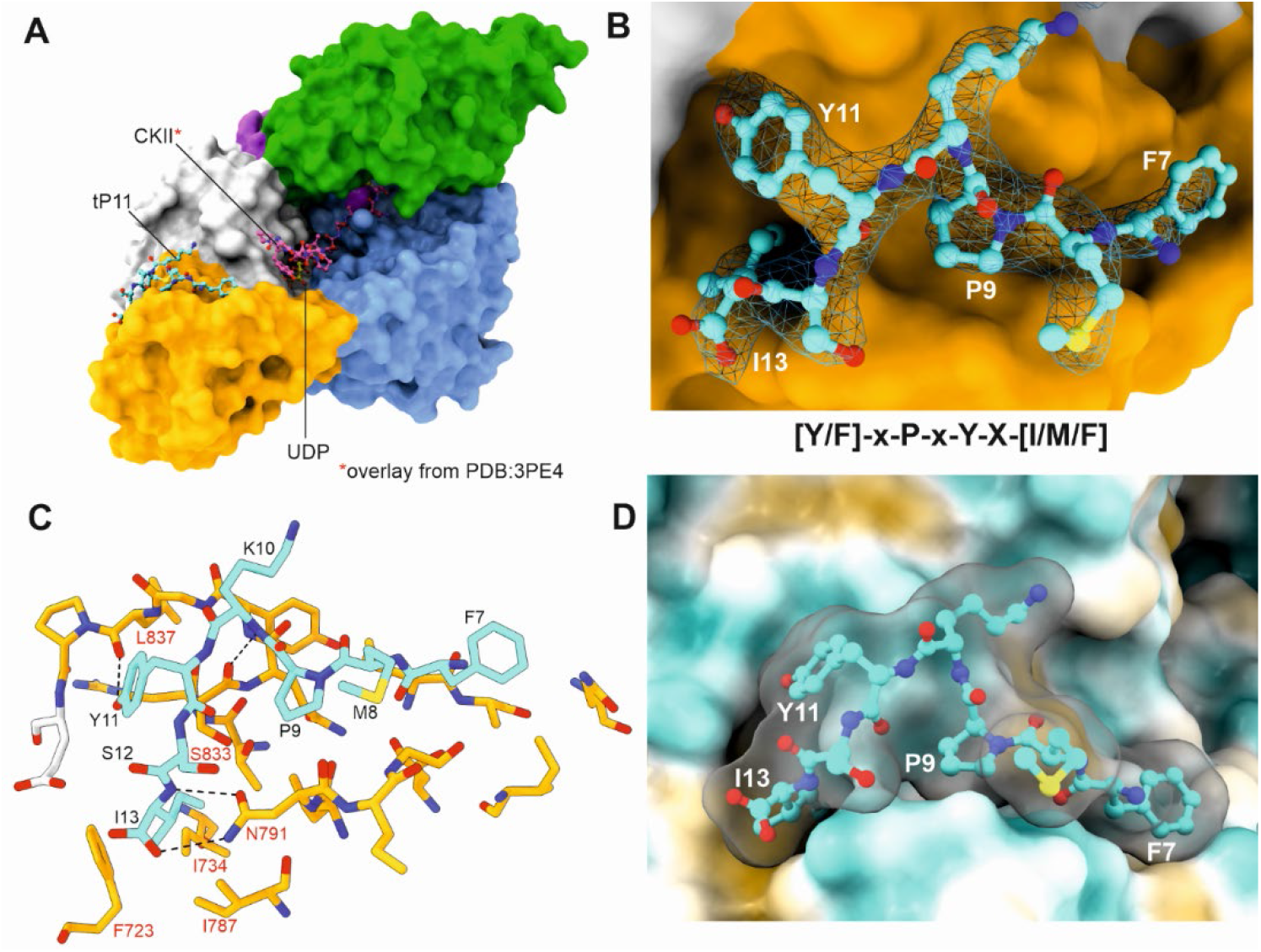
Structure of OGT in complex with tP11: (A), OGT4.5 depicted as a surface with bound tP11 peptide and UDP depicted in ball and sticks. CKII from PDB: 3PE4 also shown to demonstrate relative position between a peptide acceptor and the bound tP11 peptide. (B) Expanded view of the tP11 peptide binding to the surface of OGT, with 2*F*_*o*_*-F*_*c*_ electron density contoured to 1 σ localised to the tP11 peptide. The density is incompatible with the OGT main chain and lies across the face of the intervening domain. (C) Hydrogen bonding interactions between the tP11 peptide and OGT displayed as dashed lines. Residue numbering in black refers to P11 residues, red numbering highlights OGT residues involved in tP11 binding. (D) Surface displaying hydrophobic (brown) and hydrophilic (blue) regions of the surface of OGT. Semi-transparent surface of tP11 peptide also visualised to highlight complementary shape of the tP11 peptide to the OGT surface.

**Figure 6.**
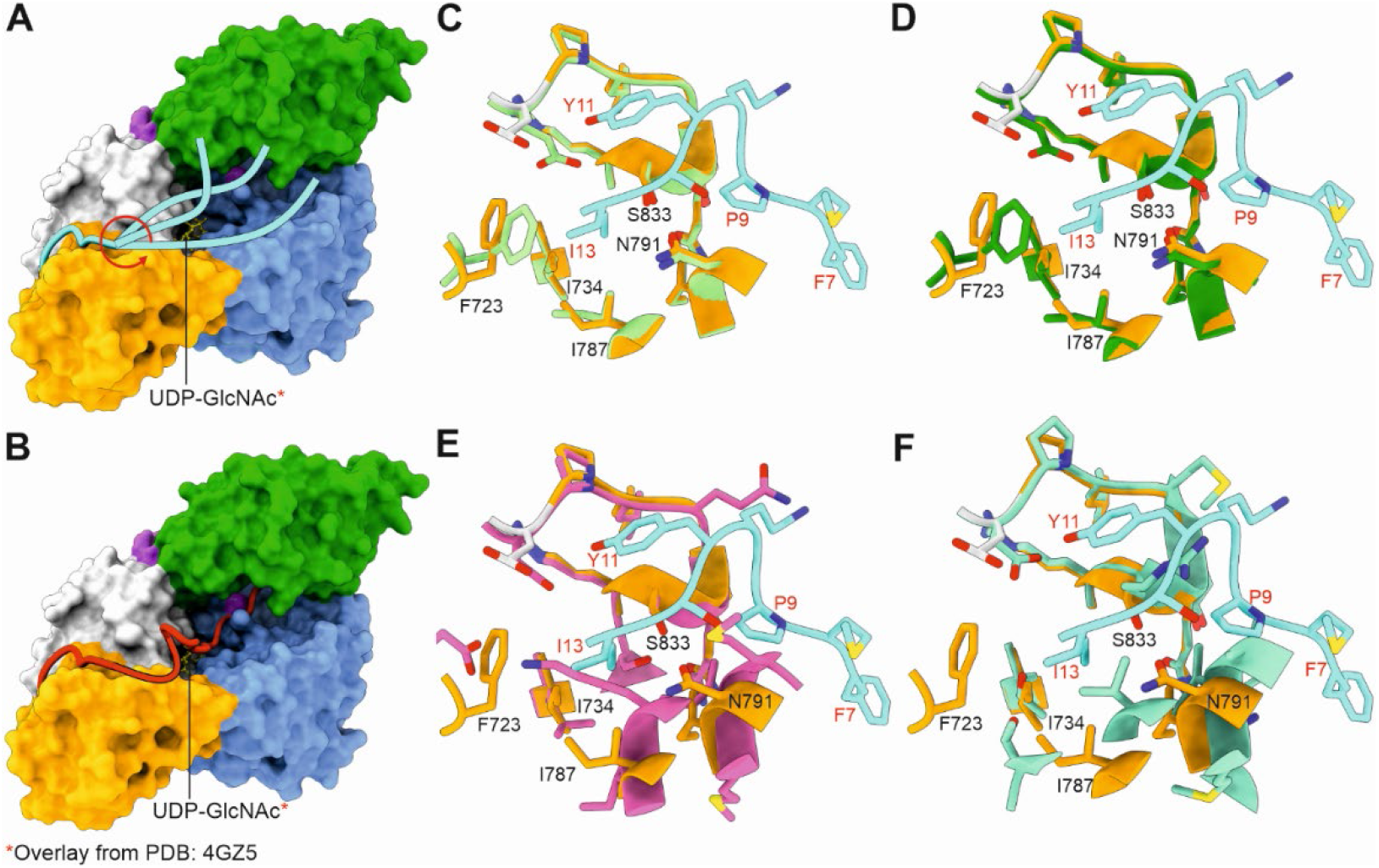
Mechanism of inhibition of OGT by exosite binding polypeptides and conservation of the OGT exosite: (A) A proposed mechanism by which peptides bearing the conserved motif prevent the binding of OGT protein substrates. (B) A second proposed mechanism by which the conserved motif in OGT protein substrates enables tighter binding and enhanced O-GlcNAcylation. (C-F) Comparisons of human OGT (*Hs*OGT) against crystal structures and AlphaFold models of OGT homologues. HsOGT superimposed onto AlphaFold generated *Mus Musculus* (C) *Danio rerio* (D) and Caenorhabditis elegans (E) OGT models. F, OGT superimposed onto the X-ray crystal structure of *Drosophila melanogaster* OGT,PDB: 5A01.(46)

Analysis of the structure of this complex revealed that residues that are not conserved within this peptide motif are solvent exposed, clearly explaining the tolerances at these sites that are seen within the deep sequencing of the pool of selected phage (**Table S1**). Conserved features of the motif can also be understood in the context of the structure. By constraining geometry, the proline residue (P9) of the peptide induces a kink that adjusts its orientation and permits its flanking residues to bind to the surface of OGT. The phenylalanine residue (F7) packs against OGT to strengthen the overall peptide interaction and may facilitate reorientation of the peptide towards the active site. These two contributions from the F7 residue provide a rationale for the loss of inhibition we observed when this residue was replaced with a proline residue in the P14 peptide. Since peptide tP11 is unable to suppress OGT activity and residues which would lie C-terminal of the consensus motif are not positioned close to the active site, we believe the disordered residues N-terminal of the consensus motif are likely to be responsible for occluding the active site and driving inhibition. Accordingly, the consensus motif likely acts as an anchoring point for the peptide, orienting these N-terminal residues so they can interfere with binding of polypeptide substrates at the active site (**Figure 6a**). This model is also consistent with the *K*_d_ value (470 ± 160 nM) for the P11 peptide being 10-fold lower than the *K*_i_ value (4.0 ± 0.9 ?M) of P11 since inhibition in this model must be exerted by a subset of the peptide conformations that occlude the active site. Such a steric occlusion model for inhibition of enzymes is, to our knowledge, unprecedented. However, though clearly different, this mode of inhibition is reminiscent of how binding of autoproteolytically processed peptides allosterically antagonize enzyme activity by binding to exosites present within certain pepsin-family aspartic acid proteases.(44)

To gain a sense for the level of conservation of this exosite among OGT orthologues, we leveraged the AlphaFold2 structure prediction database(45) to see whether OGT from model organisms that contain the Int-D domain are likely to have an exosite capable of accommodating a peptide bearing our motif. Unsurprisingly, given the high sequence similarity, the AlphaFold generated mouse OGT exosite is identical to the human, as was that of the AlphaFold OGT structure from zebrafish (*Danio rerio*) (**Figure 6c**). In comparison the AlphaFold generated structure of *Caenorhabditis elegans* and the crystal structure of *Drosophila melanogaster* have distinct differences in the features of the exosite. An arginine residue in *D. melanogaster* which replaces S833, causes significant steric hinderance to binding of the tyrosine residue of the peptide motif. In addition, residue N791 of human OGT is absent from both models. Furthermore, the hydrophobic pocket used to accommodate the I13 residue of the peptide motif has been disrupted in both organisms. These results suggest that the binding of this peptide may be a recent evolutionary feature. However, it is also possible that these organisms use their Int-D domain for other purposes yet to be defined or that other peptide sequence motifs are bound by this site in these other organisms.

Taken together, these results define an exosite on the surface of OGT that can bind peptides containing the sequence motif identified in our phage display studies. Given the strong consensus motif of peptide sequences that emerged through the selections, along with the conservation of this exosite within higher vertebrates, coupled with tolerance to variation of the residues at intervening positions, we speculate that native proteins containing this exosite-binding motif are likely present within mammalian proteomes. Moreover, the low micromolar to nanomolar binding and inhibition constants of these exosite-binding peptides are consistent with proteins containing such sequences having biological activity within cellular settings. Given their potential for disruption of the glycosyltransferase activity of OGT as we have seen here, binding of peptides and proteins to this exosite may have regulatory significance, either as we see here through inhibition of OGT but, alternatively, by serving to recruit specific protein substrates for modification (**Figure 6b**), which may either than diffuse away if binding with modest affinity or remain to antagonize OGT activity.

Given these considerations, we examined the prevalence of this consensus motif within the human proteome we conducted a bioinformatic analysis of the human proteome using the MOTIF search tool.(47) Using the consensus sequences as search string ([Y/F]-x-P-x-Y-x-[I/M/F]) that we found that 112 human genes encode proteins bearing some variation of this sequence (**Spreadsheet S1**). Of these, we curated the sequence to include only those proteins localized to the nucleus or cytoplasm, which are the cellular compartments in which OGT acts, and identified 52 proteins (**Spreadsheet S1**). Among these 52 proteins are represented protein families of various function, including transcription factors, kinases, deacetylases, and nucleic acid editing enzymes. Notably, only 9 of the 52 proteins have been previously identified as substrates of OGT that can be modified with O-GlcNAc. We expect that some proteins among this set likely bind to OGT through the identified motif and some of these may function as regulatory partners. Given the limited number of O-GlcNAcylated proteins observed in this set we reason that binding of putative partners to this exosite likely mediates inhibitory regulation of OGT. However, a number of factors will need to be considered in advancing on this question since the accessibility of the peptide motif within the protein and the secondary structure in which the motif is found are likely to influence binding to OGT. Nevertheless, additional experiments to investigate these putative interaction partners in greater detail are therefore warranted.

## DISCUSSION

The *O*-GlcNAc post-translational modification has been firmly established as a key regulator of several signalling pathways that govern important cellular processes. However, the molecular mechanisms that control the glycosyltransferase activity and substrate specificity of OGT have been slow to emerge. Indeed, the large number of *O*-GlcNAcylated targets in the human proteome raises questions about how OGT activity is regulated and how this enzyme identifies and discriminates between its great number of protein substrates. Attempts have been made to identify a generalized consensus motif surrounding serine or threonine O-GlcNAcylation sites.(8, 41, 42) Although a clear consensus motif likely does not exist, certain preferences in primary peptide sequence have been identified and these are consistent with the structural requirements of the active site of OGT.(42) The roles of the TPR region in recruiting substrates to OGT for glycosylation are gradually emerging.(22, 23, 48) Other regulatory mechanisms are also being defined,(20, 26) including the general concept that binding partners of OGT may influence substrate recruitment.(21, 24, 49) The molecular details of these interactions, however, remain unknown. Binding to the TPR region is one logical potential means of controlling substrate specificity. However, it is notable that panning of our diverse phage library did not return sequences binding to the TPR region and instead converged to deliver a sequence motif that binds to a defined exosite on the surface of the Int-D of OGT that is proximal to the active site. Notably, the amino acid residues that define the topological features of this exosite are strikingly conserved among mammals (**Figure S6**), suggesting it likely fulfills important evolutionarily conserved roles in modulating OGT function. While this exosite may either serve to recruit OGT substrates we find it is more likely that this site serves a regulatory role by recruiting OGT polypeptides, including proteins, that contain this motif such that their binding occludes the active site of OGT and prevents its subsequent binding of protein substrates. With respect to the linear peptides identified here, their low to submicromolar binding affinity suggest these peptides could be purposed in useful ways. For example, by inserting or fusing this motif to proteins one may be able to control or direct OGT activity within cells, which has emerged as a topic of some interest.(50, 51) Furthermore, chemical refinement of these structures may also offer up new chemical tools to control OGT activity in an allosteric manner as well as to interrogate the biological effects of antagonizing this exosite within cells. Perhaps most immediately salient, is that the exosite that we have uncovered is likely to have biological significance by functioning as a regulatory site for OGT at which binding of proteins and polypeptides containing this sequence motif modulate its activity. Although the short peptides identified in this study display fair but not exceptional binding affinity towards OGT, it is likely that larger proteins in cells that are capable of binding to this site may do so with significantly higher potency by their greater steric bulk, which would more effectively occlude the active site. Moreover, full length proteins may have features enabling additional contacts within the active site or elsewhere on the surface of OGT. Intriguingly, the conservation of tyrosine within this motif suggests a molecular mechanism for relief of inhibition of OGT. The sequence of this motif is compatible with kinases acting to induce phosphorylation of this tyrosine residue, which would abrogate binding of potential exosite-binding protein antagonists of OGT as the phosphate would prohibit binding to the exosite of OGT. Studies directed toward identifying protein regulatory factors and their potential control by tyrosine kinases are likely to prove a promising line of inquiry into a new mechanism for regulation of OGT activity within cells.

Notably, after our disclosure of the initial identification of this motif by phage display,^34^ and during the course of preparation of this manuscript, a pre-print describing related research appeared.^60^ That work from the Jiang group describes a closely related motif along with biophysical and structural data relating to its binding to the same site on the Int-D domain of OGT and is broadly consistent with our findings. While that work proposes this site is principally for recruitment of substrates, whereas our findings are interpreted as this site acting in an inhibitory capacity, the convergence of these two separate lines of research and similarity of core findings provides strong support for the conclusion that this exosite is most likely to play important roles in the regulation of OGT.

## MATERIALS AND METHODS

### Selection of phage peptides bound to OGT

Four rounds of panning were performed using the X15 library and histidine-tagged O-GlcNAc transferase (OGT) as a target. Manipulations of solutions and magnetic beads were performed manually in the first round; for subsequent rounds a Kingfisher Duo Prime magnetic bead purification system was used (Thermofisher). The X15 library was depleted for bead-binding phage by incubation with Dynabeads (Thermofisher) before panning. All bead incubations were set up in 1.7 ml Eppendorf tubes on a rotator. One hundred microliters of beads were washed in HN buffer (50mM HEPES and 500 mM NaCl, pH 7.4), resuspended in a total volume of 100 μL of HN containing X15 library (1.4^11^ PFU/mL), and incubated at 4°C overnight. The depleted library supernatant was then separated from magnetically immobilized beads and saved for panning. OGT-beads were prepared by incubating 10 μg of OGT protein with 20 μL of HN washed Dynabeads in total volume of 100 μL HN at 4°C overnight. OGT-beads were blocked with 2% bovine serum albumin (BSA) in HN at room temperature for 1 hour. Four replicate library panning tubes were set up, each containing depleted X15 library (10^11^ PFU/mL) with blocked OGT-beads in 1 mL of 2% BSA/HN. The tubes were incubated at room temperature for 1.5 hours. After incubation, the OGT-beads were immobilized by magnet and the supernatant was removed. The OGT-beads with bound phage were washed three times with 0.1% Tween/HN. Phage were released by incubating beads in 200 μL of 0.2M Glycine-HCl pH 2.2 on a rotator for 9 min. The supernatant containing released phage was removed from the beads and neutralized with 30 μL of 1M Tris-HCl pH 9.1. This pool of released phage was separated into three aliquots: (1) 200 μL of the pool was added to 25 mL of early log phase *E*.*coli* TG1 pRLA4 and amplified in a shaking incubator at 37°C for 4.5 hours; (2) 20 μL was used as a source of DNA for next generation sequencing; (3) the remaining phage was quantified by plaque assay. Phage amplification and subsequent polyethylene glycol (PEG) precipitation was performed as described.(52) PEG precipitated phage was resuspended, titered, and used as the library for the next round of panning. For rounds 2, 3, and 4 of panning, beads were washed 1, 3, and 3 times respectively with the Kingfisher instrument. For rounds 3 and 4 of panning, parallel negative panning controls were performed using depleted X15 libraries and beads containing no target protein. Focus library panning was performed twice, once under the same conditions as round 3 of regular panning without a bead depletion step, and the second time with an additional 3 washes, both with the addition of parallel negative panning controls.

### Biolayer Interferometry (BLI)

Biolayer interferometry experiments were performed using an Octet RED96e instrument (FortéBio, Sartorius) and 96 well black microplate (part no. 655209, Greiner). The binding assay was carried out at 25°C in assay buffer 50 mM Hepes NaOH pH 7.4, 500 mM NaCl, 1 mM TCEP, 0.1 % (w/v) BSA, 0.02 % (v/v) Tween-20 and 5 % (v/v) DMSO with sample plate shaking speed 1000 rpm. The Octet® streptavidin (SA) dip and read biosensor (part no 18-5019, Sartorius) was equilibrated in the assay buffer for 120 seconds. Then the ligand, biotinylated peptide P11 (sequence: GDFPVMFMPKYSIIF with N-terminal modification Biotin-Ahx), at a concentration 0.85 μg·mL^−1^ (400 nM) was immobilized on the biosensor for 600 seconds. The biosensor was then washed in the assay buffer for 120 seconds and dipped in the sample solution, OGT4.5 dimer, at a concentration range of 25, 125, 250, 500, 1000, and 2000 nM. Samples at a concentration of 2000 nM or buffer alone 0 nM were added in the rest of the wells to check for non-specific binding and apply correction for baseline drift. Following the association step, dissociation was performed for 600 seconds. The data correction and analysis were carried out using Octet Data Analysis High Throughput (HT) software version 12. The rate constants were determined by global fit type using all the six sample concentrations.

### X-ray crystallography

A reaction of 52.5 μM OGT4.5 with 1050 μM UDP in size-exclusion buffer was incubated for 1 hour at 4°C. The combined OGT4.5/UDP mix was then diluted 28.3 times in size-exclusion buffer containing tP11 at 57 μM to yield a final reaction mix of 1.85 μM protein, 37 μM UDP and 55 μM tP11. The OGT4.5/UDP/tP11 mixture was incubated together for 4 hours before being concentrated to ∼8 mg/ml using a 3000 MWCO Vivaspin concentrator (protein concentration determined via A_280_ using the extinction coefficient and molecular weight of OGT4.5 only). The best crystals obtained were in 1.5 M potassium phosphate dibasic, 5 % xylitol and 10 mM EDTA. Crystal drops were set up at a 2:1 ratio. Crystals were cryo-protected in mother liquor supplemented with 27% xylitol and flash cooled. Diffraction data were collected at Diamond Light Source in Oxford, UK on beamline I03. Data reduction and processing was completed through *xia2* using *DIALS* and *AIMLESS*(53, 54) A structure solution was obtained through *Phaser* using chain A of PDB 3PE3(19) as a search model. Simulated annealing was run on the structure solution through *Phenix* before cycles of refinement and interactive model building were undertaken using *REFMAC5* and *Coot*.(55-58) Residues have been numbered relative to OGT isoform 3. Figures were prepared through UCSF ChimeraX.(59)

## ACKNOWLEDGEMENTS

The authors are grateful for support from GlycoNet, the Canadian Glycomics Network (CD-1) and the Canadian Institutes of Health Research (PJT-156202). Natural Sciences and Engineering Council of Canada (RGPIN-05426). We thank Diamond Light Source for access to beamline I03 (proposal number mx24948). GJD thanks the Royal Society for the Ken Murray Research Professorship and RWM for the associated PDRA funding (RP\EA\180016). We also wish to acknowledge Dr. Johan Turkenburg and Sam Hart for helping coordinate data collection. DJV thanks the Canada Research Chairs program for support as a Tier I Canada Research Chair in Chemical Biology. The authors also thank the Centre for High-Throughput Chemical Biology (HTCB) for access to core facilities.

## Notes

https://www.rcsb.org/

