## Supporting Materials for "Phage display uncovers a sequence motif that drives polypeptide binding to a conserved regulatory exosite of O-GlcNAc transferase"

**This PDF file includes:**

Supporting text

Figures S1 to S7

Tables S1 to S6

Legends for Datasets S1

SI References

**Other supporting materials for this manuscript include the following:**

Datasets S1

### SUPPORTING METHODS

*OGT expression and purification*

The plasmid containing the gene encoding full-length human OGT in a previously-reported pET28a vector(1) was transformed into competent *Escherichia coli* (*E.coli*) BL21 (DE3) cells (Invitrogen). Successful transformants were cultured in Terrific Broth supplemented with 50 µg/mL kanamycin at 37°C until an $OD_{600}$of 1.8 was reached. Protein expression was induced with 0.2 mM isopropyl β-D-thiogalactoside at 16°C for 18 h. Cells were the harvested and resuspended in 2 mL BugBuster protein extraction reagent (EMD Millipore) per gram of cell pellet in the presence of 1 mg/mL lysozyme, 0.2 mg/mL DNase and an EDTA-free protease inhibitor tablet (Roche). The mixture was gently rocked at 4°C for 30 min and then clarified by centrifugation for 2 x 30 min at 18,000 x *g*. The cell lysate was then applied to a 1 mL HisTrap nickel column (GE Healthcare) which was pre-equilibrated in 50 mM HEPES, 300 mM NaCl, 20 mM imidazole and 1 mM DTT at pH 7.5. The protein was purified using an imidazole gradient of 20-500 mM over 50 min on an AKTA FPLC (GE Healthcare). Fractions judged as pure were pooled and dialyzed at 4°C in 50 mM HEPES buffer containing 500 mM NaCl and 1 mM DTT at pH 7.4. Aliquots of OGT were flash frozen and stored at -80°C for future use.

Briefly, OGT4.5 bearing a 3C-protease cleavable N-terminal 6xHisTag was expressed in *E.coli* BL21(DE3) cells. Cultures were grown to an $OD_{600}$ of 1.0-1.2 in TB media, at 37 °C, in the presence of 35 µg/ml Kanamycin. Cells were chilled at 16 °C for 30 mins before being induced with 0.2 mM IPTG. Cultures were shaken overnight at 16 °C before being harvested at 5500 *g* for 20 mins. Pellets were stored at -20 °C until use. To purify OGT, pellets were resuspended in buffer A (20 mM Tris pH 7.4, 150 mM NaCl, and 40 mM imidazole), lysozyme, DNAse I, and 1 mM AEBSF and then lysed by cell disruption. Insoluble material was removed by centrifugation at 41700 g for 1 hour at 4 °C. The soluble fraction was loaded onto a 5 ml HisTrap FF column (GE Healthcare) equilibrated in buffer A. Non-specific interactors were removed by 10 column volumes of buffer A before OGT4.5 was eluted by a stepwise gradient of buffer B (20 mM Tris 7.4, 150 mM NaCl, and 400 mM imidazole) in buffer A. OGT4.5 containing fractions were pooled and supplemented with THP to a final concentration of 1mM. Samples were then incubated with HRV 3C protease at a ratio of 1 µg protease to 100 µg OGT4.5 overnight at 4 °C. The protein solution was diluted to 15 mM NaCl using 20 mM Tris pH7.4 before application to a 5 ml HisTrap FF column (GE Healthcare) in tandem with a 5 ml HiTrap Q HP column (GE Healthcare) equilibrated in buffer C (20 mM Tris and 40 mM imidazole, pH 8). Non-specific contaminants were removed with 10 column volumes of buffer C. OGT4.5 was then eluted with a stepwise gradient of 20 mM Tris pH 8 through to 20 mM Tris pH 8, 1 M NaCl. Fractions containing pure protein were pooled and concentrated using a 10K MWCO Vivaspin® spin concentrator (Sartorius) to 2 ml. Protein was then size-excluded on a 16/600 Superdex 200 column (GE Healthcare) in 20 mM Tris pH 8 and 150 mM NaCl. Fractions containing pure protein were pooled, supplemented with THP to 1 mM and concentrated as before. Protein was quantified by $A_{280}$ and then snap frozen and stored at -70 °C till use.

*X15 Phage library design*

The 48HD X15 (X15) library is a phage peptide library constructed in the phage vector M13KE, an M13mp19 derivative that can be propagated without the need for helper phage superinfection.(2) The library was constructed by inserting a synthetic oligonucleotide library (Twist Bioscience, San Francisco) into the vector via KpnI and EagI sites as described previously.(3) The resulting library gives rise to phage encoding pIII protein with 15 random amino acids immediately downstream of the leader peptidase cleavage site in pIII. The low valency of pIII proteins on M13 (5 copies) gives rise to libraries that are well suited for discovery of high affinity ligands. A second type of library, called a 'focus library' was created by choosing enriched sequences obtained after panning as a new pool of oligonucleotides to be cloned into the modified M13KE vector. Details on the choice of sequences for that library are described in the Sequencing/Bioinformatics section below.

*Sequencing of Phage DNA*

Aliquots of phage pools taken at various stages during panning were used as a source of DNA for next generation sequencing on the Illumina NextSeq platform (Illumina, San Diego). These phage pools were converted to Illumina-compatible short dsDNA fragments by PCR using primers with four-nucleotide-long barcodes to trace multiple samples in one Illumina run as previously described.(4) A full analysis of the Illumina sequencing results from all rounds was performed. The results suggested that the output of round 3 had converged to a smaller set of sequences, which were potential binders. Meanwhile, the OGT output of round 4 (**Figure S6**) showed modest enrichment compared to round 4 Input. This indicated that round 4 input saturated with OGT binders. Therefore, we focused on the analysis on the data sets collected from round 3.

Differential Enrichment (DE) analysis was applied on the data sets collected from round 3 (**Table S1**): 1) Before depletion ($R3^{PreDep}$), 2) Input ($R3^{Input}$), 3) OGT output ($R3^{OGT}$), and 4) Ni-NTA Beads output ($R3^{Ni\text{-}NTA}$). First, singleton sequences, which were observed only once through all replicates in the dataset, were filtered out. Then, the Trimmed Mean of M-values (TMM) normalization and a negative binomial model were applied to train the data sets (5), which estimated the normalized counts per million (CPM) of each sequence in each dataset, the fold change (FC) and its corresponding p-value. Out of all the sequences observed in round 3, 645 sequences satisfying the criteria: 1) FC $R3^{OGT}/R3^{Input}$ and $R3^{OGT}/R3^{Ni\text{-}NTA}$ are greater than 2 with p-value less than 0.05; and 2) FC $R3^{Input}/R3^{PreDep}$ is above 0.5. The second criteria ensured that the sequences were not potential bead binder, which were supposed to be removed or decreased through the depletion process. These 645 sequences were encoded by using the BLOSUM62 matrix (**Table S3**), then the density-based spatial clustering of applications with noise (DBSCAN) algorithm was applied to classify those sequences into 28 clusters.

Then, considering 2 factors; i) whether the sequence was observed in round 4 OGT output, and ii) whether the sequence contains the most abundant motif pattern after a motif analysis on the 645 enriched sequences, 23 sequences (**Table S4**) were selected to build the focus library. Due to the capacity of library generation, each of the 23 X15 sequences were first truncated to 5 X11 sequences. For example, the sequence NQYQPQYQILYANNR resulted in sequences NQYQPQYQILY, QYQPQYQILYA, YQPQYQILYAN, QPQYQILYANN, and PQYQILYANNR. Therefore, 115 X11 sequences were first included in the focus library. Then 7 (**Table S5**) out of the 115 sequences were selected for full positional maturation, replacing each position with the other 19 amino acids. The resulting sequences were included in the focus library as well.

A single-round panning without depletion was performed on the focus library thereafter (**Figure S7**). Three data sets were collected in this focus panning: 1) input ($FL^{In}$), 2) OGT output ($FL^{OGT}$), and 3) Ni-NTA beads output ($FL^{Ni\text{-}NTA}$). A similar DE analysis was applied to these data sets as well to estimate the normalized counts per million (CPM) of each sequence in each data set. FC ratios $FL^{OGT}/FL^{In}$ and $FL^{OGT}/FL^{Ni\text{-}NTA}$ were used to select the sequences enriched most for the further study.

*In vitro OGT activity assay*

Activity assays were performed as previously described.(6) Briefly, a master mix containing various concentrations of UDP-GlcN-BODIPY and biotinylated peptide acceptor is made up in PBS containing 12.5 mM $MgCl_2$ and 1 mM DTT at pH 7.4. The reaction is commenced by the addition of recombinant OGT in PBS to a final well volume of 25 µL with a final OGT concentration of 20-200 nM. The plate is then incubated at ambient temperature for up to 60 min; during which time the reaction rate was shown to be linear. The reaction is terminated by the addition of 25 µL stop mix containing 2 mM UDP and 0.2 mg/mL streptavidin-coated magnetic beads (Trilink Technologies M-1002). The plate is incubated at room temperature for at least 30 min to allow for streptavidin-biotin binding, and is then subjected to an automated washing procedure using a BioTek EL406 plate washer containing a magnetic adapter and a BioTek 384F magnet (12 wash cycles, 100 µL PBS dispensed per well, 6 mm aspiration height offset, 4 min initial resting time on magnet, 1 min rest between washes). At the end of the wash cycle, a final dispense of 50 µL PBS per well is performed using the syringe dispenser of the BioTek washer. The fluorescence signal is read using a Biotek Neo2 multimode plate reader using 490 nm excitation and 525 nm emission wavelengths. Each well is read using an area scan function which records the mean signal of 9 individual scans in a grid pattern with 1.0 micron spacing between each point. Data were analyzed and plotted using GraphPad Prism 5.

*OGT inhibition assays*

$IC_{50}$ experiments were carried out using the general procedure for the OGT activity assay, with the following modifications. Inhibitor stocks in DMSO were diluted to various concentrations and were pre-incubated with recombinant OGT on ice for at least 15 minutes. The reaction was commenced by addition of a master mix containing both 3 µM UDP-GlcN-BODIPY and 10 µM Biotin-HCF peptide, corresponding to their respective $K_M$ values. Reactions proceeded at room temperature for < 60 min, during which the reaction progress was shown to be linear. The relative activity of OGT in the presence of inhibitor was assessed by normalizing fluorescent signal against controls which contained only DMSO (100% activity) and controls which contained 2 mM UDP (0% activity). Percent activities were plotted as a function of $\log_{10}$ concentration of inhibitor. Curves were fitted using a sigmoidal 4-parameter log(inhibitor) vs response function in GraphPad Prism 5 [$Y=100/(1+10^{((LogIC50-X)*HillSlope))}$]; $IC_{50}$ values were calculated from the inflection point of sigmoidal curves. Full $K_i$ inhibition experiments were carried out in a similar manner, except that the concentration of substrate was varied to at least 5-fold above and 5-fold below the $K_{M\,(app)}$ in the presence of varying concentrations of inhibitor. $K_i$ values were calculated using the non-linear regression function of GraphPad Prism 5.

### SUPPORTING FIGURES

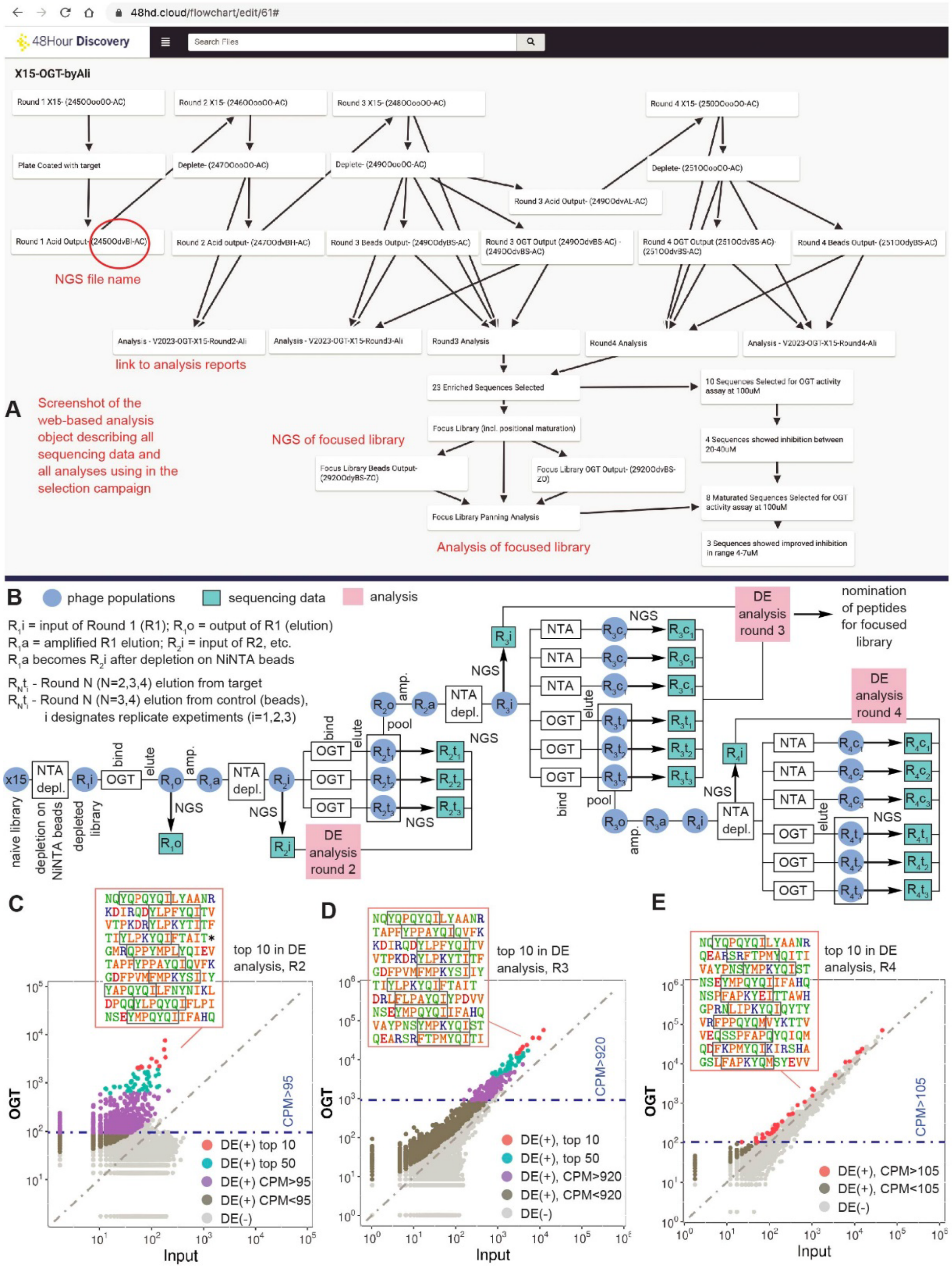


**Figure S1 – Outline of phage display screening approach and sequence conservation of the peptide binders of OGT:** (A) Web-based **flow chart object** describing all panning and next generation sequencing (NGS) steps performed in the selection. Each box leads to an NGS data set or analysis reports. Boxes that contain a 12-character alphanumeric IDs can be used to locate the corresponding NGS data. For example 245OOdvBI-AC (circled red) is the ID of the NGS analysis of the first round of panning of X15 naïve library against OGT. Boxes labeled “analysis” contain populations of sequences generated by differential analysis of NGS samples and analysis reports. Link to the flow chart and all sequencing datasets can be shared upon request. (B) Schematic of four rounds of the phage panning procedure using bead-immobilized OGT. In each round, the phage library was depleted against Ni-NTA beads to generate the input library, which was incubated with bead-immobilized OGT, followed by washing and elution. Samples from the resulting phage library were submitted for next generation sequencing (NGS), and also amplified in *E.coli* for the next round screening.  The blue circles identify samples of phage at various steps in the procedure and the green squares indicates steps at which next-generation sequencing was performed. Various combinations of sequencing samples were used to preform differential enrichment analysis (red boxes). (C-E) Bioinformatic analysis of Round 2, 3 and 4 of screening NGS data. Each dot represents a sequence that observed in the NGS data, color coding of dots is provided in the legend. Axes represent normalized counts per million (CPM) for NGS dataset obtained from input and selection on OGT. Controls datasets are not shown for clarity, but they can be seen in the full analysis reports (SI). Top 10 sequences that showed significant enrichment in OGT samples over input or beads are shown for each dataset. Grey boxes are highlighting the (YF)-x-P-x-Y-x-(IML) 7-mer amino acid motif

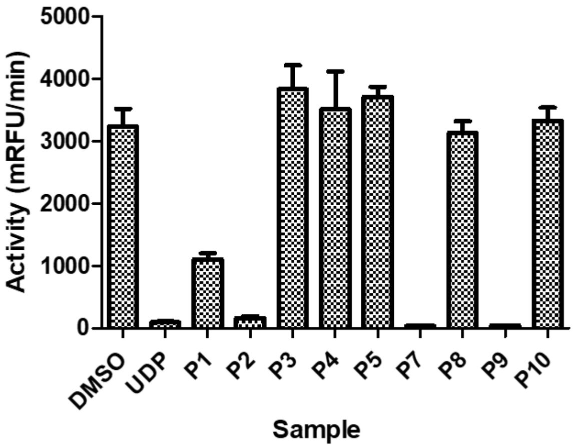


**Figure S2 - Inhibition of prioritized peptides identified from phage panning towards OGT:** Peptides were incubated with OGT at a concentration of 100 µM and relative glycosylation of the HCF-Serine peptide acceptor was determined using a fluorescence-based activity assay. Peptides displaying greather than 50% inhibition of OGT at this concentration were prioritized for follow-up experiments. Error bars represent S.E.M. of triplicate measurements.

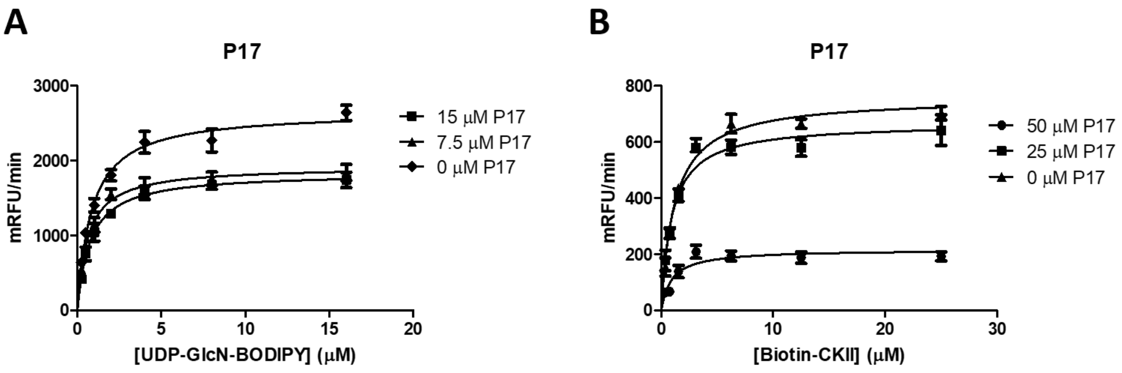


**Figure S3 - Phage peptide P17 inhibits OGT non-competitively with respect to both UDP-GlcNAc and CKII peptide acceptor substrate:** (A) Michaelis-Menten curves of UDP-GlcN-BODIPY in the presence of varying concentrations of peptide P17. (B) Michaelis-Menten curves of Biotin-CKII in the presence of varying concentrations of peptide P17. Increasing concentrations of P17 result in a reduction of $V_{max}$ without significantly affecting $K_M$. Data processing and curve fitting were performed using Graphpad Prism 5.

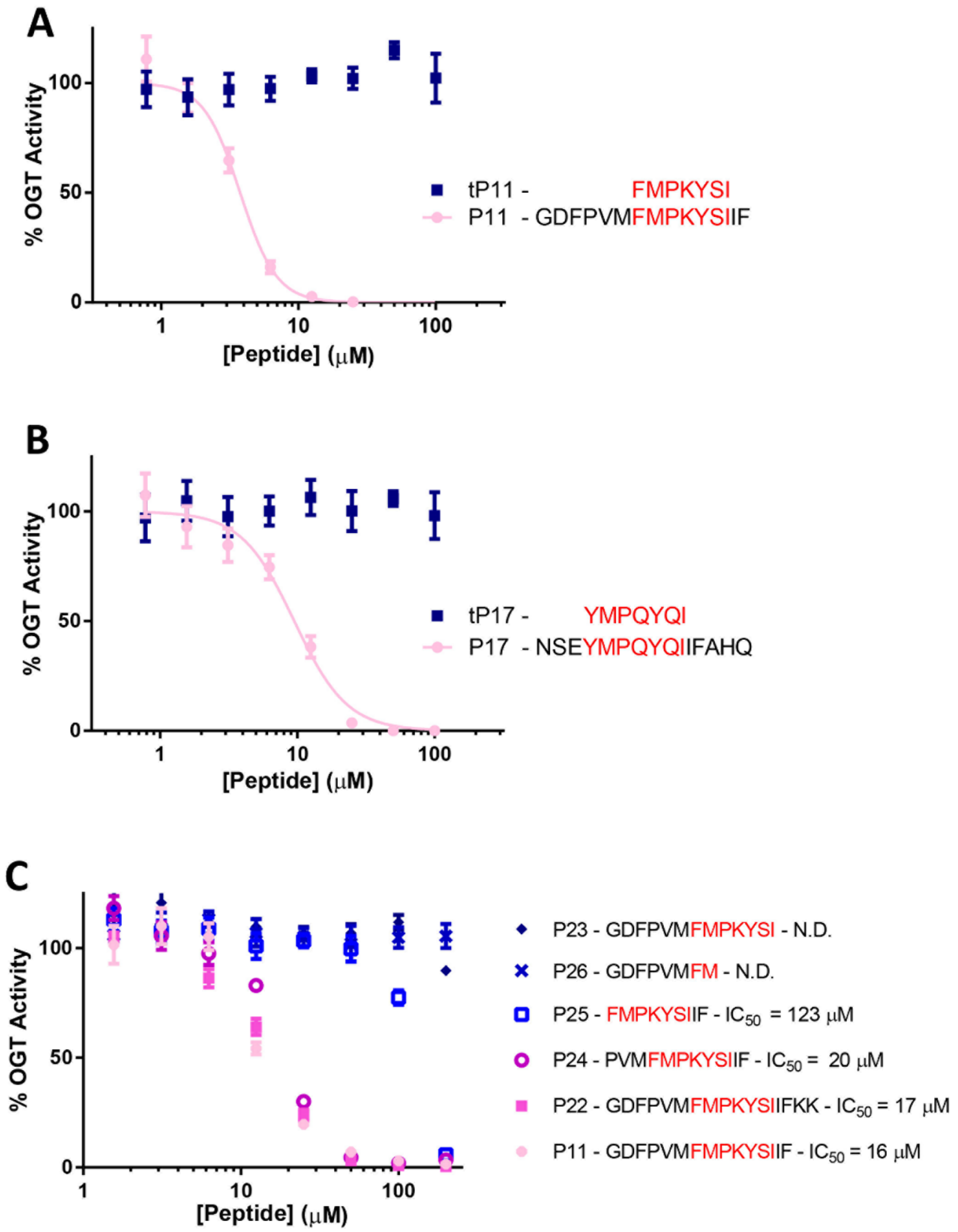


**Figure S4 - Dose-dependent inhibition of OGT activity by modified phage peptides:** (A) Concentration-response curves of P11 and the truncated analogue (tP11) comprising just the amino acids spanning the conserved motif. (B) Concentration-response curves of P17 and the truncated analogue (tP17) comprising just the amino acids spanning the conserved motif. (C) Concentration-response curves of truncated and modified peptides. The consensus motif determined from analysis of phage panning results is shown in red for each peptide. % OGT activity was normalized relative to controls containing 1% v/v DMSO (100% activity) or 1 mM UDP (0% activity). All assays were performed in buffer containing 0.02% Triton X-100 detergent. Sigmoidal curves were fit using a four-parameter log(inhibitor) vs response formula in GraphPad Prism 8. Error bars represent S.E.M. of quadruplicate measurements.

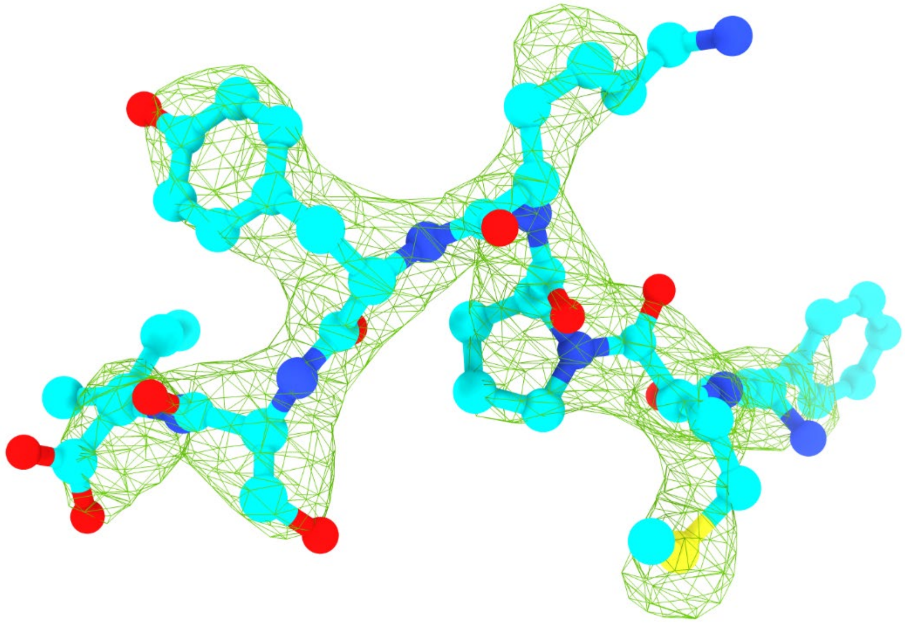

**Figure S5. Omit $F_o$-$F_c$ electron density of the tP11 peptide (chain E).** Electron density displayed in green mesh at 3 σ and carved at 1.9 Å around the peptide shown in ball and stick. Carbon atoms are shown in cyan, oxygen atoms in red, nitrogen in blue, and sulfur in yellow.

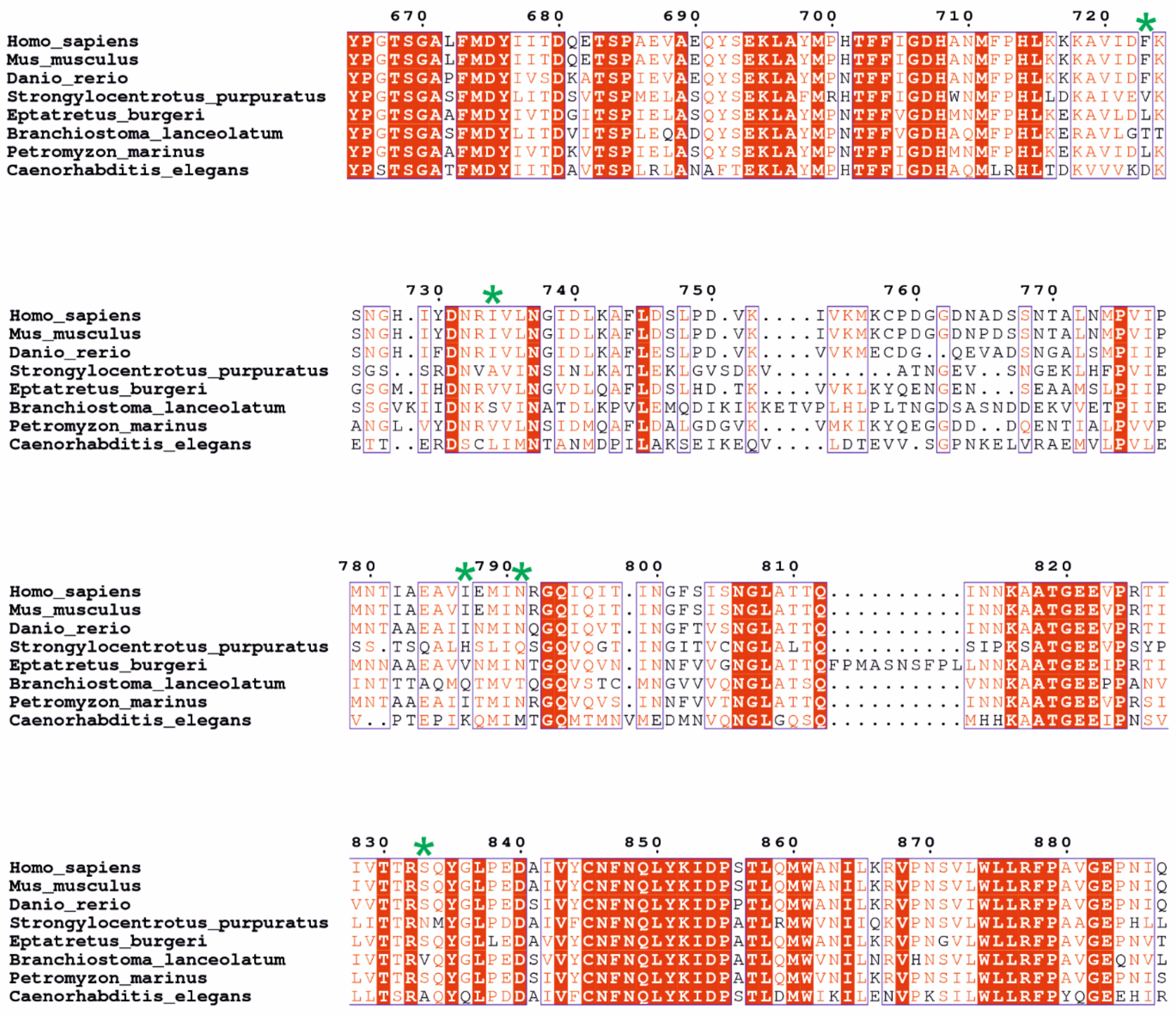


**Figure S6 - Sequence alignment of OGT homologues from different organisms:** UNIPROT IDs for the proteins are as follows: *Homo sapiens* (O15294-1), *Mus musculus* (Q8CGY8), *Danio rerio* (B3DKP3), *Strongylocentrotus purpuratus* (A0A7M7NIJ2), *Eptatretus burgeri* (A0A8C4Q3M1), *Branchiostoma lanceolatum* (A0A8K0A0K7), *Petromyzon marinus* (S4RUA6), *Caenorhabditis elegans* (O18158).

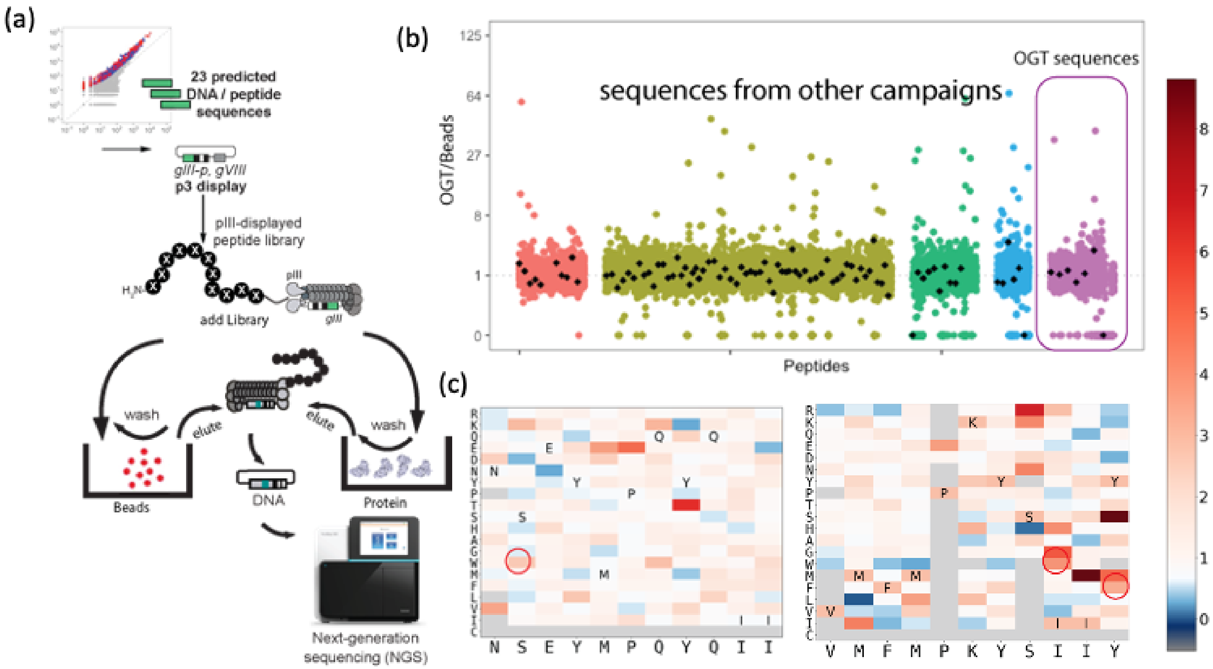


**Figure S7** – Focus library screening. (a) Schematic of focus library screening procedure using bead-immobilized OGT. Sequences selected from OGT screening and other 4 campaigns against other targets were included in a focus library containing in total 12,000 sequences. DNA oligonucleotides of those 12,000 sequences were cloned into pIII-displayed phage library and then screened with both NiNTA beads and OGT protein. (B) Manhattan plot of focus library screening results, OGT/Beads represent the FC $FL^{OGT}/FL^{Ni\text{-}NTA}$. (C) Heatmap shows FC $FL^{OGT}/FL^{Ni\text{-}NTA}$ of variants from full position maturation of two sequences NSEYMPQYQII and VMFMPKYSIIY. The variants highlighted in red circles were selected to derive the three peptides in Figure 3b and other sequences were rejected based on various criteria (lack of significance of the observed enrichment, low copy number in the sequencing sample, etc.)

**SUPPORTING TABLES**

**Table S1:** List of next-generation sequencing data with links to the openly available datasets.

| DataSet | Round 1 | Round 2 | Round 3 | Round 4 |
|---|---|---|---|---|
| **Before Depletion** | | 20190221-246OOooOO-AC | 20190221-248OOooOO-AC | 20190221-250OOooOO-AC |
| **Input** | 20190221-245OOooOO-AC | 20190221-247OOooOO-AC | 20190221-249OOooOO-AC | 20190221-251OOooOO-AC |
| **OGT** | 20190221-245OOdvBI-AC | 20190221-247OOdvBH-AC | 20190221-249OOdvBS-AC | 20190221-251OOdvBS-AC |
| **Ni-NTA Beads** | | | 20190221-249OOdyBS-AC | 20190221-251OOdyBS-AC |

**Table S2.** Sequences and inhibition values of truncated and extended peptides

| Peptide ID | Sequence | OGT $IC_{50}$ (µM) |
|---|---|---|
| P11 | GDFPVM**FMPKYSI**IF | 6.0 ± 0.4 |
| tP11 | **FMPKYSI** | > 200 |
| P21 | KKGDFPVM**FMPKYSI**IF | 5.3 ± 0.4 |
| P22 | GDFPVM**FMPKYSI**IFKK | 16 ± 4 |
| P23 | GDFPVM**FMPKYSI** | > 200 |
| P24 | PVM**FMPKYSI**IF | 20 ± 3 |
| P25 | **FMPKYSI**IF | 120 ± 14 |
| P26 | GDFPVM**FM** | > 200 |
| P17 | NWE**YMPQYQI**IFAHQ | 5.1 ± 0.6 |
| tP17 | **YMPQYQI** | > 200 |

**Table S3.** Data collection and refinement statistics for OGT4.5 in complex with UDP and P11s. Values in parenthesis are for highest-resolution shell. Data was derived from a single crystal.

| | OGT4.5-tP11 complex |
|---|---|
| Accession code | 8CM9 |
| **Data Collection** | |
| Spacegroup | *P*321 |
| Cell dimensions | |
| a, b, c (Å) | 272.97, 272.97, 142.6 |
| α, β, γ (°) | 90, 90, 120 |
| Resolution (Å) | 122.1 - 2.8 |
| $R_{meas}$ | 0.286 (2.65) |
| $R_{p.i.m}$ | 0.062 (0.563) |
| $I/\sigma I$ | 9.3 (1.4) |
| $CC_{1/2}$ | 0.996 (0.619) |
| Completeness | 100 (100) |
| Multiplicity | 21.2 (22) |
| **Refinement** | |
| Resolution | 122.4 – 2.8 |
| No. of reflections | 142053 |
| $R_{work}/R_{free}$ | 0.196 / 0.224 |
| No. of atoms | |
| Protein | 21,858 |
| Ligands (UDP + tP11) | 344 |
| Water | 236 |
| Average *B*-factors | |
| Protein | 63.3 |
| Ligands | 79.7 |
| Waters | 44.4 |
| RMSDs | |
| Bond angles (°) | 1.544 |
| Bond lengths (Å) | 0.0089 |

**Table S4: BLOSUM 62 Matrix**

| Amino Acid | Feature Vector |
|---|---|
| A | [0.077, -0.916, 0.526, 0.004, 0.240, 0.190, 0.656, -0.047, 1.357, 0.333] |
| R | [1.014, 0.189, -0.860, -0.609, 1.277, 0.195, 0.661, 0.175, -0.219, -0.520] |
| N | [1.511, 0.215, -0.046, 1.009, 0.120, 0.834, -0.033, -0.570, -1.200, -0.139] |
| D | [1.551, 0.005, 0.323, 0.493, -0.991, 0.010, -1.615, 0.526, -0.150, -0.282] |
| C | [-1.084, -1.112, 1.562, 0.814, 1.828, -1.048, -0.742, 0.379, -0.121, -0.102] |
| Q | [1.094, 0.296, -0.871, -0.718, 0.500, -0.080, -0.442, 0.202, 0.384, 0.667], |
| E | [1.477, 0.229, -0.670, -0.355, -0.284, -0.075, -1.014, 0.363, 0.769, 0.298] |
| G | [0.849, 0.174, 1.726, 0.093, -0.548, 1.186, 1.213, 0.874, 0.009, 0.242] |
| H | [0.716, 1.548, -0.802, 1.547, 0.350, -0.785, 0.655, -0.076, -0.186, 0.990] |
| I | [-1.462, -1.126, -0.761, 0.382, -0.599, 0.276, -0.132, 0.198, -0.216, 0.207] |
| L | [-1.406, -0.856, -0.879, -0.172, 0.032, 0.344, 0.109, 0.146, -0.436, -0.021] |
| K | [1.135, -0.039, -0.802, -0.849, 0.819, 0.097, 0.213, 0.129, 0.176, -0.850] |
| M | [-0.963, -0.585, -0.972, -0.528, 0.236, 0.365, 0.062, 0.208, -0.560, 0.361] |
| F | [-1.619, 1.007, -0.311, 0.623, -0.549, 0.290, -0.021, 0.098, 0.433, -1.288] |
| P | [0.883, -0.675, 0.382, -0.869, -1.243, -2.023, 0.845, -0.352, -0.421, -0.298] |
| S | [0.844, -0.448, 0.423, 0.317, 0.200, 0.541, 0.009, -0.797, 0.624, -0.129] |
| T | [0.188, -0.733, 0.178, -0.012, 0.022, 0.378, -0.304, -1.958, 0.149, 0.063] |
| W | [-1.577, 2.281, 1.166, -1.610, 0.122, 0.239, -0.542, -0.398, -0.349, 0.499] |
| Y | [-1.142, 1.740, -0.582, 0.747, -0.119, -0.475, 0.241, -0.251, 0.713, -0.251] |
| V | [-1.127, -1.227, -0.633, 0.064, -0.596, 0.158, 0.014, 0.016, 0.251, 0.607] |

**Table S5:** 23 Sequences selected to build the focused peptide library.

| | | |
|---|---|---|
| TAPFYPPAYQIQVFK | YDKVMPEYQIKYTTS | QVFWPQYQINIPKLN |
| NQYQPQYQILYAANR | GPRNLIPKYQIQYTY | FPSFLPAYPIEYSNT |
| NSEYMPQYQIIFAHQ | VAYPNSYMPKYQIST | GIVNKYLPKYQIETI |
| TMPRDFTPKYQIMVK | APQQYLPAYQIHYNK | VRFPPQYQMVYKTTV |
| DRLFLPAYQIYPDVV | VTPKDRYLPKYTITF | QEARSRFTPMYQITI |
| GDFPVMFMPKYSIIY | KDIRQDYLPFYQITV | VEQSSPFAPQYQIQM |
| ETYFPPVYQMEFVQR | SPFMPMYPIVPNRSD | QSLVTQYYPKYQIRI |
| NVRFPPMYQMVWTSR | QWDYYPPAYQIRTSV | |

**Table S6:** The seven peptide sequences selected for full positional maturation.

| | | | |
|---|---|---|---|
| TAPFYPPAYQI | YPPAYQIQVFK | NQYQPQYQILY | NSEYMPQYQII |
| EYMPQYQIIFA | VMFMPKYSIIY | PNSYMPKYQIS | |
